# Ouvrai: Opening access to remote VR studies of human behavioral neuroscience

**DOI:** 10.1101/2023.05.23.542017

**Authors:** Evan Cesanek, Sabyasachi Shivkumar, James N. Ingram, Daniel M. Wolpert

**Affiliations:** Zuckerman Mind Brain Behavior Institute, Department of Neuroscience, Columbia University, NY, USA

## Abstract

Modern virtual reality (VR) devices offer 6 degree-of-freedom kinematic data with high spatial and temporal resolution, making them powerful tools for research on sensorimotor and cognitive functions. We introduce Ouvrai, an open-source solution that facilitates the design and execution of remote VR studies, capitalizing on the surge in VR headset ownership. This tool allows researchers to develop sophisticated experiments using cutting-edge web technologies like the WebXR Device API for browser-based VR, without compromising on experimental design. Ouvrai’s features include easy installation, intuitive JavaScript templates, a component library managing front- and back-end processes, and a streamlined workflow. It also integrates APIs for Firebase, Prolific, and Amazon Mechanical Turk and provides data processing utilities for analysis. Unlike other tools, Ouvrai remains free, with researchers managing their web hosting and cloud database via personal Firebase accounts. Through three distinct motor learning experiments, we confirm Ouvrai’s efficiency and viability for conducting remote VR studies.

## Introduction

Virtual reality (VR) gives neuroscientists the ability to create immersive 3D environments and record the movements of research participants at high spatial and temporal resolution. By robustly engaging the sensorimotor system to create a feeling of presence, VR provides a higher degree of ecological validity than standard computer-based laboratory methods. At the same time, VR offers fine-grained control of stimulus properties, task sequencing, and other aspects of experimental design, unlike observational research. For instance, motor control researchers often use VR to perturb the relation between the visual and actual configuration of the body during reaching movements. Here we will focus on applications of VR in the study of human movement, but the range of potential research applications is extremely broad. For example, cognitive neuroscientists have used VR in a laboratory setting to study brain mechanisms related to locomotion, navigation, spatial cognition, object manipulation, multisensory integration, threat detection, theory of mind, and more, while clinically-focused researchers have studied potential therapeutic applications of VR for psychiatric disorders, pain management, and rehabilitation ^1–3^.

To date, nearly all VR studies have been moderated laboratory studies. Participants are recruited from the local community and schedule a visit to a physical laboratory, where an experimenter personally guides them through the experiment. This is a reliable process, but it can be frustratingly slow and costly. Crowdsourced research has emerged as an effective way to bypass this data collection bottleneck, and “online studies” surged in popularity during the COVID-19 pandemic, especially among neuroscientists ^4^. Crowdsourcing hubs such as Prolific and Amazon Mechanical Turk (MTurk) give researchers access to a world-wide population of participants—provided that their experiments can be accessed through a web browser.^i^

Some researchers have attempted browser-based studies of human motor learning that rely on standard desktop computer peripherals ^5–13^. Unfortunately, this approach has significant limitations. Due to the properties of different input and display devices, the mapping between real movements of the hand and the digitized movements recorded by web browsers is nonlinear and variable. A quick twitch of the finger on a trackpad can produce the same data as a 10-cm mouse movement involving the entire arm. Conversely, two identical movements performed with different trackpads (or mice) can produce completely different data. Additionally, operating systems and browsers may unpredictably throttle the rate at which mouse or trackpad events are processed. Moreover, using a mouse or trackpad to hit visual targets on a 2D screen already entails a context-specific realignment of visual and bodily coordinates which is uncharacteristic of natural movements. These issues complicate data analysis and provoke reasonable concerns about how “motor” these kinds of tasks really are, casting doubt on the generality of findings. Fortunately, VR provides a potential solution to all of these problems by displaying visual stimuli and recording movement data in well-aligned 3D coordinate frames at a high, stable refresh rate, producing consistent data even across different devices.

Although total sales figures for VR headsets (including offerings from Meta, HTC, Sony, HP, Pico, Huawei, and more) tend to be a trade secret, it is nonetheless clear that current ownership and usage levels are sufficient to support crowdsourced research. By one conservative estimate, over 11 million Meta Quest 2 headsets were sold by the end of 2022, with prices as low as $300 ^14^. Meta executives have said that the number is close to 20 million ^15^. Meanwhile, a survey of 98,000 U.S. adults found that 23% of respondents own or have used a VR headset and, among owners, 80% reported using it once a month or more ^16, 17^. Of course, sales and usage should not be conflated with availability of research participants. A more pertinent test of participant availability can be conducted using Prolific’s study setup tool. In early 2023, we found that applying the “VR headset ownership” screener yielded 9,297 matching participants, approximately 8% of Prolific’s pool of active participants in the past 90 days. In the three studies reported here, which had median (per-participant) completion times of about 18 minutes, we collected data from 10 participants per experiment within 90 minutes of posting the experiments online. Larger-sample studies may achieve even faster rates of data collection, as there are more “open slots” available at once.

In developing Ouvrai, our goal was to make it easy for researchers to conduct remote VR studies with crowdsourced participants. As we will show, remote VR studies allow researchers to parallelize data collection from many participants in experiments with immersion, interactivity, and data quality rivaling what is currently achievable in a physical laboratory. Our work builds on recent engineering successes in both VR and web standards. The latest generation of standalone VR headsets feature six-degree-of-freedom (6-DoF) tracking of the head and handheld controllers (i.e., position and orientation) at 120 Hz, which is sufficient even for studying rapid reflex responses. Some devices also track the hands, providing kinematic data for 25 rigid bodies (19 joints) per hand. The WebXR Device API specifies how custom, device-agnostic VR applications (apps) can be run entirely within a VR headset’s web browser ^18^. Alongside the development of WebXR, JavaScript packages for 3D graphics such as Three.js have made it easy to write code that creates interactive virtual environments. Despite these advances, the barrier to entry for a neuroscientist interested in using these technologies remains high. There is a clear need for software that provides a low-friction entry into remote VR research through a streamlined development and deployment workflow.

Although a handful of software solutions exist that aim to help researchers design online studies, these suffer from a variety of drawbacks. First, to our knowledge, no existing solution provides support for VR or even for 3D graphics (see the Discussion for detailed explanation of why the Unity game engine is not a good solution for online studies). Second, most solutions are not end-to-end, but merely libraries or graphical interfaces that aim to help researchers create experiments (e.g., jsPsych, lab.js, and PsychoPy), often by emphasizing low- or no-code experiment building which tends to limit flexibility. Third, all existing end-to-end solutions—those that provide hosting, database, and crowdsourcing integration, like Qualtrics, Gorilla, Pavlovia, and Open Lab—charge annual or per-participant fees. In our view, these three drawbacks are symptoms of the same underlying problem: existing solutions oversimplify the process of running remote studies by hiding or removing opportunities for experimental control and study management. Meanwhile, we have observed that most of our colleagues and trainees in neuroscience would prefer to exercise a higher degree of control over their experiments. They also tend to be comfortable writing code to define their design, stimuli, and procedures, or are willing to learn. Thus, in developing a VR-enabled, end-to-end, and free remote research workflow, we generally opted to leave experimental control and study management capabilities in the hands of the researcher, while keeping the process simple enough for researchers with only basic knowledge of scripting and the command line. This balance is achieved through a carefully crafted study pipeline and powerful core components that provide an easy user experience for simple experiments, but make it straightforward to ramp up complexity as needed.

To understand what is possible with Ouvrai, we recommend that readers go to https://ouvrai.com to try the VR demo experiments, which are shortened versions of the experiments reported below. For those without a VR headset, there are also demos of computer-based and tablet-based experiments. Readers are then encouraged to install and set up Ouvrai to get a feel for how it works. Examining the JavaScript code for the demos in conjunction with the descriptions in the Methods below should provide a thorough introduction to Ouvrai, allowing researchers to get started coding and running their own experiments.

In the Results section, we first explain the design goals for Ouvrai and how these guided the software engineering process. We begin by explaining the remote study infrastructure, followed by the layout of the code that researchers must write to create an experiment with Ouvrai. We then describe the experimental results of the three motor learning validation studies. The experimental methods are described in the Methods section at the end of the article. Note that the following is intended as an overview of Ouvrai; detailed documentation geared toward practical use is available at https://ouvrai.com.

## Results

### Remote study infrastructure

The first design goal of Ouvrai was to assemble the infrastructure required to run and manage a remote study: web hosting, a cloud database, and crowdsourced recruitment. Fig. 1 shows a schematic of the Ouvrai infrastructure, including these resources, which are represented within the tan block on the right (hosting and database) and the light blue block near the bottom (crowdsourcing). These are hosted on the web, separate from the researcher’s computer (large green block), but they are accessible using Ouvrai commands (bold green text on arrows connected to the central Ouvrai diamond).

**Fig 1.**
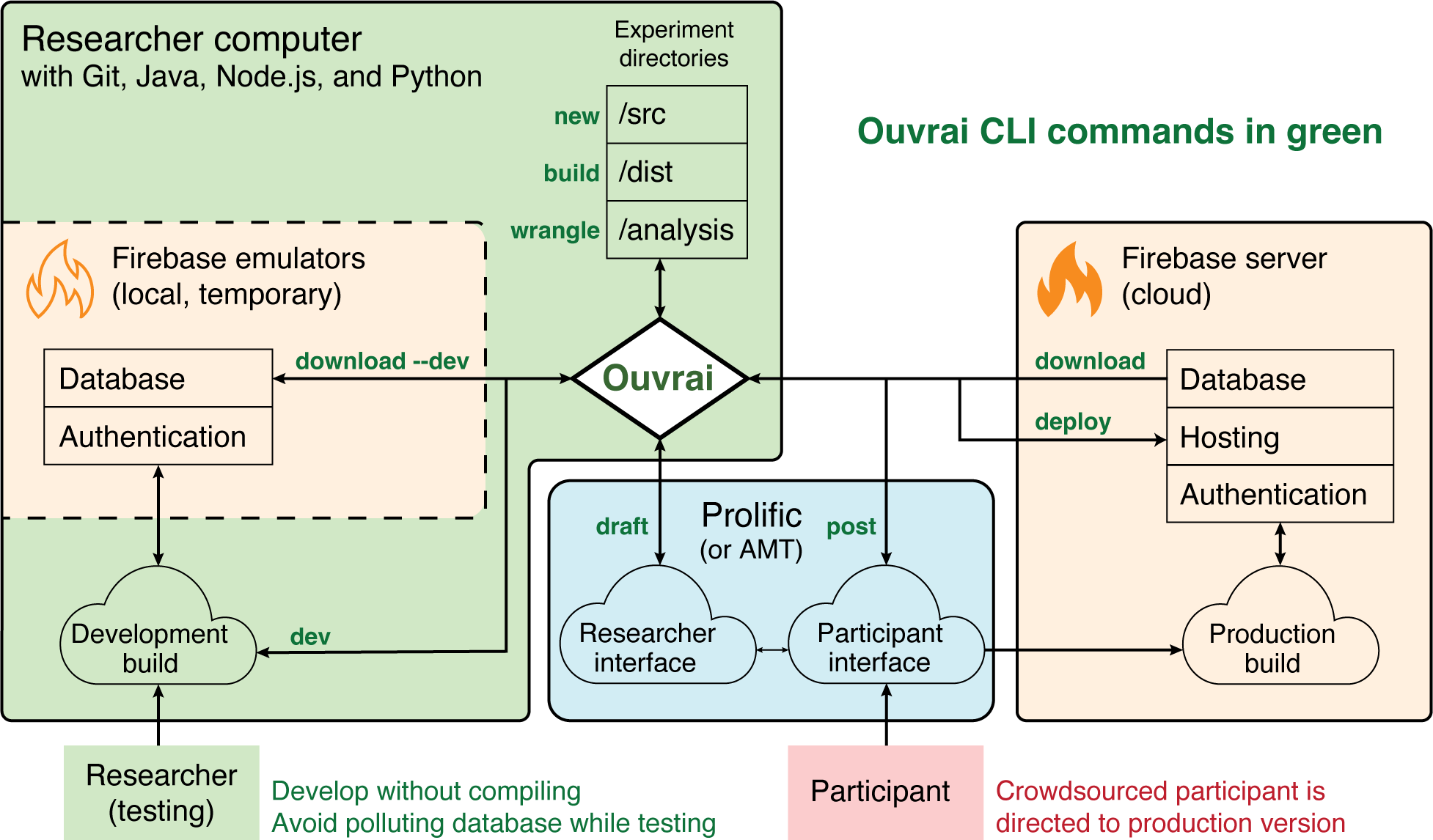
Schematic of Ouvrai infrastructure. The large green block represents the researcher’s computer. The diamond-shaped Ouvrai node (center) represents the command-line interface (CLI) used to manage studies in Ouvrai. Each Ouvrai study is housed within an experiment directory that includes several subfolders (*src, dist, analysis*, etc). An Ouvrai study is essentially a web app written in JavaScript (mainly in the *src* subfolder) and designed to be publicly hosted on Google Firebase (orange block, right). Firebase provides free cloud database, hosting, and authentication services, which are critical to online studies. The final “production” version of a study (from the *dist* subfolder) is *deployed* to a Firebase hosting URL (e.g., https://cognitivescience.web.app). When opened in a web browser, the study communicates with the Firebase servers to (1) authenticate participants by assigning them anonymous IDs and (2) write study data to a secure database, which can be *downloaded* by the researcher (into the *analysis* subfolder). Before *deploying* the production version, however, the researcher will spend some time developing their study. This process is streamlined by Ouvrai’s *dev* mode, which creates a locally hosted, development version of the study that the researcher can test in a web browser on their own device (green rectangle, bottom-left). While in *dev* mode, changes to the study code are immediately reflected in the browser without the need to compile or restart the local server. This local development server is powered by Vite, an open-source build tool. In parallel, Firebase database and authentication services are locally emulated by the Firebase Emulator Suite (orange dashed region, left), working just as they would in production, but without cluttering the production database with test data. Ouvrai makes it easy to *download* data from the emulated database and wrangle this data into a set of data tables to be analyzed (see Table 1 for details). When the study code is finalized, the researcher *builds* the production version and *deploys* to Firebase. Finally, the researcher drafts a post to appear on a crowdsourcing platform (blue block) that advertises the study to participants and, when ready to make the study public, *posts* it. At this point, participants who are logged in to the crowdsourcing platform (red rectangle, bottom) will see the post and, if they accept a spot in the study, get a link to the Firebase web hosting site where the production version has been *deployed*.

**Table 1.** The four data tables exported from Ouvrai studies. Rows are in descending order of temporal resolution. Data from different tables can be easily merged, grouped, and otherwise manipulated as needed in downstream analyses.

| Data type | Update rate | Description of data table |
| --- | --- | --- |
| Frame | 60–120 Hz | Each row is one render frame. Contains data sampled at the device refresh rate, such as position, orientation, and millisecond timestamps. In VR, refresh rates are typically 90–120 Hz. Desktop and mobile refresh rates are typically 60 Hz, but dropped frames are common in practice. |
| State | Several times per trial (~5 Hz) | Each row is one transition of the finite state machine that controls the flow of the experiment. Contains state names and timestamps, as well as any other data sampled at state transitions. For example, in the VR experiments reported here, we sample the head position and orientation only at state transitions in order to reduce the required Firebase database storage capacity, as instantaneous head position was not relevant to our study. |
| Trial | Once/trial | Each row is one trial of the experiment. Contains parameters and data that can change from trial to trial, such as the location of a target or the participant's reaction time. |
| Participant | Once/experiment | Each row is one participant. Contains parameters and data that do not change over the experiment such as the location of a fixed home position or the participant's demographic information. |

#### A. Hosting and Database

For the hosting and database requirements, we required a free, secure, and easy-to-use cloud service, as most researchers do not have the resources, time, or skills to set up and maintain their own web servers. We chose to use Firebase, a Google product with a free tier. On the free tier, each Firebase project is granted one database instance with 1 GB of storage space and supporting up to 200 simultaneous connec-tions ^19^. Given that 120-Hz, 6-DoF kinematic data from the head, hands, and possibly additional sources (e.g., all the joints on both hands) accumulate rapidly over time (*∼*3 MB/min in informal testing), we recommend regularly downloading data from the Firebase cloud database, storing it permanently elsewhere, and clearing the database to avoid reaching the limit. By default, each Firebase project is associated with one Hosting site where one experiment (i.e., web app) can be deployed at a time. The URL of this site is based on the name of the project. For instance, the default site URL for a project named Ouvrai would be https://ouvrai.web.app. Additional Hosting sites with different URLs can be created within a Firebase project, allowing multiple experiments to be deployed simultaneously. In Fig. 1, these cloud database and hosting resources are depicted inside the tan block on the right side, separate from the researcher’s computer (large green block). Note that the other tan block contained within the researcher’s computer represents the Firebase Emulator Suite. This is an additional Firebase utility involved in Ouvrai’s development workflow, allowing researchers to quickly and easily test their experiments during development by locally emulating the behavior of Firebase cloud services. This is advantageous because researchers can develop and test an experiment without having to repeatedly build and publicly deploy the production version and without polluting their limited-storage cloud database.

The functions that save data to Firebase during an Ouvrai study are configured to follow a specific database structure. The database structure is depicted in detail in Supp. Fig. 1, but researchers do not need to understand this structure, as Ouvrai provides easy access to extract the formatted data. An accompanying set of database security rules enforce this structure while preventing unauthorized database reading and writing. These security rules depend on the Anonymous Authentication feature of Firebase (the third Firebase service depicted in Fig. 1). After setting up Firebase during the Ouvrai installation process, researchers can immediately get started collecting data from the template experiments without any additional configuration. The Ouvrai Command Line Interface (CLI; central diamond in Fig. 1) provides commands (green bold text) that allow researchers to interact with Firebase, for example to deploy an experiment to web hosting so it is publicly accessible, or to download the participant data associated with each experiment.

Alternative products, including Microsoft Azure and Amazon Web Services (AWS), offer web hosting and cloud database services and include free tiers. However, at present they require users to switch to a payas-you-go plan after 12 months. After this time, although basic web hosting and cloud database services may remain free under certain usage limits, the researcher becomes vulnerable to unexpected charges if they accidentally exceed these limits. This is not a concern with Firebase, unless the researcher knowingly opts in to the pay-as-you-go plan, which involves providing credit card information.

#### B. Crowdsourced recruitment

Once a study is deployed to Firebase web hosting, the researcher must provide the study link to participants. Crowdsourcing services allow researchers to post these links to a large pool of remote participants and pay compensation to those who complete the study. We aimed to provide a unified interface for posting studies and reviewing submissions on Prolific and MTurk, two of the most popular crowdsourcing services. The pros and cons of crowdsourced data collection on different platforms have been discussed extensively elsewhere ^20, 21^.

When posting a new study, configuration options including the title, description, reward amount, estimated completion time, and desired sample size should be specified by editing the study configuration file located within the relevant Ouvrai experiment directory on the researcher’s computer (this file is not explicitly shown in Fig. 1, but it would be located alongside the directories displayed at the top of the large green block). The study configuration file also allows the researcher to specify an allowlist and/or blocklist to restrict participation. These lists can be specified in terms of specific participants and/or previous experiments. Then, the Ouvrai CLI can be used to post a draft version of the study for the researcher to review and test on the web interface of each service, and to publicly post the finalized version to the participant pool (see *draft* and *post* commands in Fig. 1, bold green text). Some researchers may prefer to use the Prolific web interface to post new studies, as it can be more intuitive and provides some additional features not available in Ouvrai. This does not interfere with the ability to use Ouvrai to manage the rest of the study (or to *draft* the post with Ouvrai, and then switch to the Prolific interface to modify and publish it). The MTurk web interface does not provide sufficient functionality to post Ouvrai studies, so we do not recommend using it (see full documentation for more details). Researchers who post a study to MTurk must use the Ouvrai dashboard (which is opened with the CLI *launch* command) to monitor which participants have completed the study and to approve or reject their submissions.

Crowdsourcing fees are the only unavoidable cost involved in using Ouvrai. Prolific and MTurk both charge a percentage fee on the compensation paid to participants. Currently, MTurk charges a 20% or 40% fee^ii^, while Prolific charges a 33% fee on the compensation provided to participants. In agreement with studies comparing ethical concerns and data quality across crowdsourcing services ^20, 21^, we currently recommend that researchers prioritize Prolific. Prolific also provides built-in screeners for VR headset ownership and many other participant characteristics, as mentioned above. These screeners would have to be implemented manually with a qualification test or a separate screener study on MTurk.

### Streamlined installation, setup, and workflow

The second design goal of Ouvrai was to minimize friction and barriers to use, starting with installation and setup, and continuing through study development and deployment, all the way to data analysis. Although we have streamlined this process to a great degree, Ouvrai still requires researchers to be comfortable navigating their computer’s file system and running basic commands in a command shell.

#### A. Quick installation and setup

Before getting started, researchers must install Git, Node.js, Java, and Python, and create a Firebase account with one Firebase project using the web interface, then install the Firebase Command Line Tools. However, after the one-time setup of these prerequisites (documented in detail at https://ouvrai.com), researchers can get started with Ouvrai by cloning the GitHub repository, opening a command shell, and running three simple commands. When the researcher runs the *setup* command, they are led through a series of prompts (most users will just press Enter at all prompts) that connect Ouvrai with the researcher’s Firebase project, set up the required resources for this Firebase project, and create template Firebase configuration files within Ouvrai. These template files will be automatically copied into each new experiment going forward (using the *new* command, Fig. 1). Thus, Ouvrai condenses many tedious steps into a single *setup* command, so the researcher does not have to spend time understanding the many different services offered by Firebase (most of which are not needed), navigating the Firebase web interface to set up project resources, or figuring out how and where to create the required local configuration files. The *setup* command also creates a Python virtual environment inside the Ouvrai installation and installs the required Python packages needed for the data pre-processing utilities (described below), thus isolating these installations from other Python environments the researcher may have on their computer. This allows the researcher to use these data utilities without ever directly reading or writing any Python code.

#### B. Development and deployment workflow

After the setup is complete, the researcher can immediately create a new experiment based on one of the included templates by running the *new* command, then start the development server with the *dev* command, which launches a locally hosted version of the experiment that will immediately reflect any changes to the code, and download their own testing data in a format that matches how the real data will be formatted using the *download* command with the --*dev* flag. When the experiment is working as desired in *dev*mode, the researcher runs the *build* command to create the bundled and minified production version of the experiment. These convenient development features rely on Ouvrai’s unique combination of the Firebase emulators with the Vite development server (Fig. 1, large green block). Finally, the *deploy* command sends the production files to a Firebase Hosting site of the researcher’s choosing, where it will be publicly available. At this point, any visitor to the site who begins participating in this study will have their data saved to the Firebase cloud database.

With Ouvrai, testing and debugging VR experiments in dev mode is easy for Android-based VR headsets, such as the Meta Quest. Simply connect the headset to the development computer with a USB cable, enable port forwarding in Google Chrome (from chrome://inspect), and navigate to the locally hosted experiment in the headset’s web browser (at http://localhost:5173 by default). For untethered development, researchers can use Android Debug Bridge (ADB) over Wi-Fi. With either approach, the researcher can debug their experiment code using the console output and other features of the Chrome Developer Tools (from chrome://inspect). Detailed documentation on development workflows for VR, computers, and mobile devices are provided at https://ouvrai.com.

#### C. Data analysis utilities

After downloading participant data (whether real data or test data), researchers can convert the unprocessed JavaScript Object Notation (JSON) data files into a set of four tidy data tables at different levels of resolution (Table 1) using Ouvrai’s data pre-processing methods (*wrangle* command, Fig. 1). These tables can be exported in a variety of formats, providing a convenient starting point for further analysis in Python, Matlab, R, or other analysis software.

#### Creating an experiment

The third design goal of Ouvrai was to provide researchers with a focused but flexible study development experience. This was a critical goal because the powerful study infrastructure and the streamlined workflow described above are only valuable if researchers using Ouvrai can manage to implement functional experiments in JavaScript.

Behavioral studies, whether lab-based or remote, have a number of requirements that must be implemented in order to be successful. These include experiment configuration (e.g., specifying global param-eters and selecting among different conditions), generation of flexible trial lists, controlling the flow of the experiment during and between trials, presenting stimuli to participants, recording responses (including kinematic and other sampled data), and saving data in a useful format. Online studies have additional requirements, including remotely obtaining informed consent and demographic information from participants, detecting and blocking users who should not attempt to complete the experiment, communicating with database and crowdsourcing services, and controlling browser and/or VR headset features such as fullscreen, pointer lock, and spatial tracking.

To keep things simple for those who are new to developing experiments as web applications, Ouvrai limits the main experiment code for each study to one file that is focused on familiar aspects of experimental design like procedure, stimuli, and trial sequence. Below, we describe the typical structure of Ouvrai experiment code and explain how these aspects are implemented. First, however, we must briefly introduce the Ouvrai component library, which handles a wide variety of critical functions (setup, front-end, and back-end) behind the scenes, in order to provide a focused development experience.

#### A. Component library

The Ouvrai component library dramatically reduces the amount of code that the researcher has to write and/or modify in the main experiment file. The library is designed as a JavaScript ES module, which means that the researcher can import the various library components in the main experiment file. Ouvrai components offer reliable solutions to many of the basic requirements of an online study so the researcher does not need to worry about them at all (though researchers are free to examine and improve the source code). Meanwhile, when it comes to procedure, stimuli, trial sequence, and other aspects that the researcher must be able to customize, Ouvrai components provide convenient interfaces with documented functions that help build the envisioned study. Importantly, all of the components were designed with the aim of avoiding any strong limitations on what a researcher might create. Lastly, since Ouvrai studies are JavaScript ES modules, researchers are not limited to importing functions and classes from the Ouvrai component library; they have access to a wide range of third-party JavaScript packages available via Node Package Manager (npm) or elsewhere, as described below (see Extensibility).

#### B. Single-file experiment code

Although an Ouvrai experiment consists of JavaScript code that is distributed across a large number of files, including the component library and third-party imports, researchers only need to manage one file per experiment. This file consists of the definition of variables and other functions, and code to control the flow of the experiment. When a participant visits the study page, their browser first waits until the HTML document structure has been fully loaded, and then it executes the “main” function defined in this file. The main function must perform six distinct tasks, which we describe in sequence below.

##### i. Experiment configuration

The core of an Ouvrai study is the Experiment component, which is instantiated at the beginning of the study code. The Experiment component instantiates a number of “nested” components that are used throughout the study, such as the Firebase component for saving data, the State component for managing experiment flow, and the SceneManager component for rendering stimuli. The Experiment component can be configured with many different options. For example, some options pertain to requirements of the study, such as requiring users to access the study from a specific device (e.g., VR headsets but not computers or mobile devices), while other options pertain to features of the study, such as choosing 2D or 3D graphics and enabling hand tracking, and other options that are helpful during study development, such as skipping the consent form and saving the trial list so that complicated designs can be examined for accuracy.

##### ii. Trial procedure

After instantiating an Experiment, the researcher must initialize the finite state machine (FSM) component. A FSM can be in exactly one of a finite number of states at any given time. The FSM can change from one state to another depending on a set or rules and the change from one state to another is called a transition. The researcher provides a list of state names as well as a function that will be called at all state transitions, typically used to record the transition with a timestamp in the saved data. The different states of the FSM control what operations are performed at different phases of a trial. The FSM also defines the conditions for transitioning to other states (including “interrupt” states that pause the experiment under certain circumstances). The details of FSM processing are specified by the researcher in a function called “stateFunc”, which will be described below. An FSM can typically be drawn as a flowchart. The FSM for a simple reaching experiment (Experiment 1, described below) is depicted in Supp. Fig. 2. For example, in an experiment there could be a DELAY state that waits 200 ms when the hand is the home position before presenting a target and transitioning to the START state which then waits for the user’s hand to exit the home position before transitioning to the REACH state.

**Fig 2.**
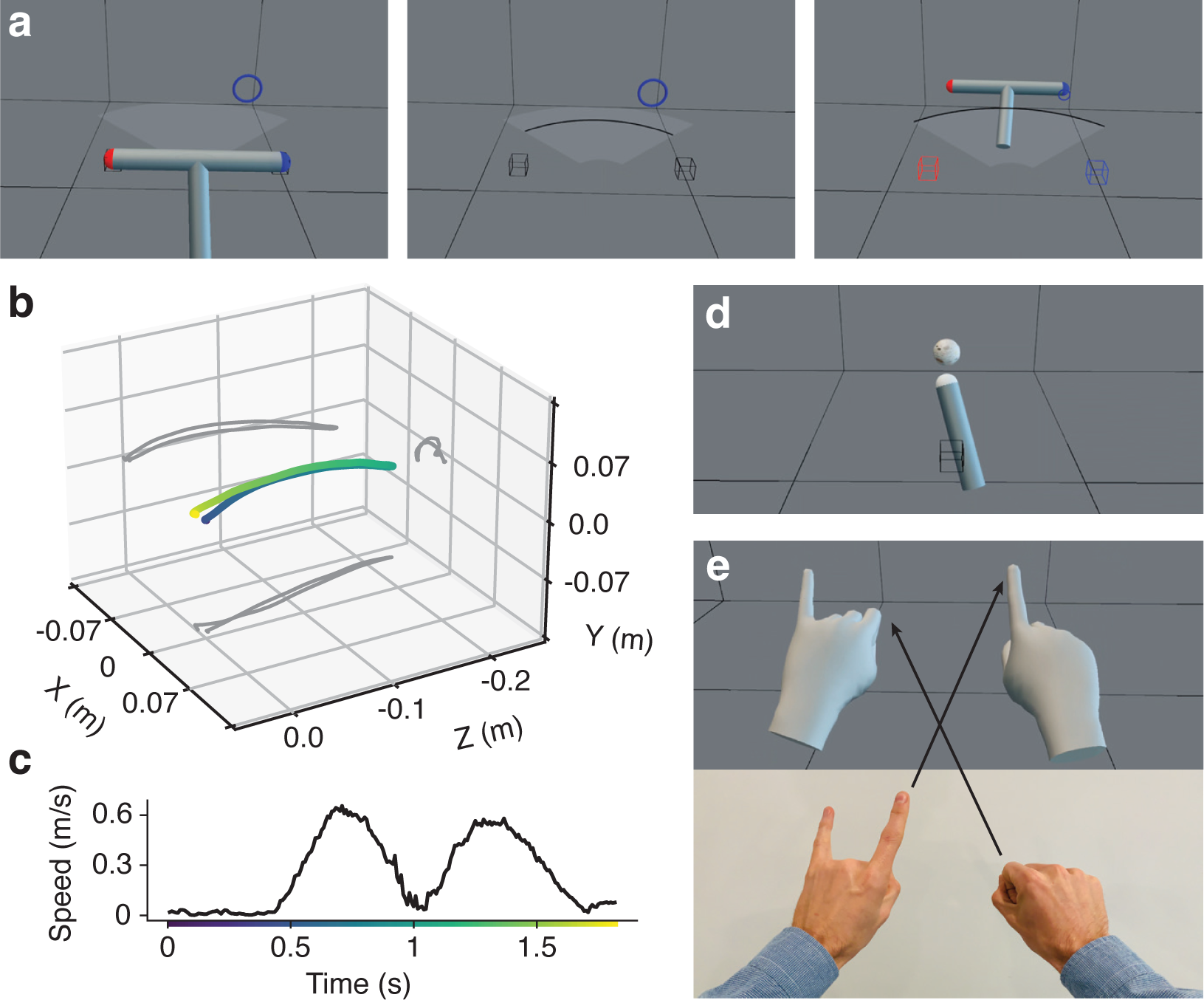
VR screenshots and representative kinematic data from Ouvrai studies. (a) Three screenshots from a single trial of Experiment 2, in which participants reached with a T-shaped tool to hit target rings with the “control point” of the same color (the red or blue end of the tool). Tool at the home position (left), in the middle of a “terminal-feedback” reach (middle), and after successfully hitting the blue target ring (right). (b) A reach path including the return movement in 3D coordinates with the origin at the home position (color represents time, see x-axis of panel c for colorbar legend). Gray shows projections of the path onto the three planes. (c) First-difference speed profile of the out-and-back reaching movement from panel (b) shows quality of the data. (d) Screenshot from Experiment 3 shows the pointer tool (also used in Experiment 1) and one distal target. (e) Screenshot from the Ouvrai demo accessing and manipulating hand tracking data (top). Photograph of the actual hand configuration at the moment of the screenshot (bottom). The demo swaps the index fingers between the two hands, so a physical movement of the right index finger produces visual movement of the left index finger in VR, and vice versa.

##### iii. Stimuli

Next, the researcher must create stimulus objects using Three.js. The display of these objects during the experiment is controlled by the “displayFunc” function, which will be described below. We chose to rely on Three.js as our web 3D engine because it is an open-source engine with first-class support for WebXR, and it has a large user community that has created a variety of different components which can be layered to include support for physics, AI, and more. The Three.js documentation explains in detail how to create stimulus objects and provides many examples. In particular, Three.js classes are useful for graphics, audio, and linear algebra (e.g., Vector3, Mesh, SphereGeometry, AudioSource etc.).

##### iv. Trial sequencing

Lastly, the researcher must create a trial sequence by instantiating one or more Block components. Each Block component corresponds to a different phase of an experiment, such as pre-exposure and exposure to a visuomotor rotation (or other experimental perturbation). To instantiate a Block component, the researcher must specify the values of variables (e.g., the direction and distance to targets) on the various trials of the block, as well as options pertaining to the entire block (e.g., the name of the block, the number of repetitions, and how to order the trials within each repetition).

##### v. Experiment loop

After these four steps have been performed, the experiment loop can be started. This loop involves three core functions (calcFunc, stateFunc, and displayFunc), which the researcher must define in the remainder of the main experiment code. These functions are executed in sequence when each frame is rendered, usually at the refresh rate of the display device.

- **calcFunc**: This function is called to process data, such as controller position. For example, it can use the controller position and the home positions to determine whether the participant is currently at the home position. This information can then be used in multiple states, allowing researchers to avoid repeating code in the stateFunc. Conversely, however, researchers should avoid doing expensive computations in calcFunc if they will only be needed in one or two states.
- **stateFunc**: This function implements the FSM, applying the rules for transitions between states (Supp. Fig. 2). It also monitors timers in order to implement time-out events (e.g., waiting for a participant to move). An FSM can be in only one state at any given time. The experiment proceeds by transitioning between states, such as ‘SETUP’, ‘START’, ‘DELAY’, ‘GO’, ‘MOVING’, etc. The function also monitors the conditions that initiate “interrupt” states (Supp. Fig. 2, inset). For example, if an experiment requires the right-hand VR controller to be connected, the FSM will transition to the ‘CONTROLLER’ state when necessary, pausing the experiment until the controller is detected.
- **displayFunc**: This function updates the graphical objects in the Three.js scene and ultimately issues the call to render to the participant’s display device. In general, scene objects should be created before starting the experiment loop, using imported Three.js classes. The displayFunc is used primarily to change the properties of objects, including controlling which objects are visible in a particular frame and setting the color or position of objects. Note that the properties of scene objects can also be changed elsewhere in the code, if it is more convenient or efficient, such as in “event handler” functions that respond to specific browser events, or in the FSM, for instance setting the target position in the ‘SETUP’ state when initiating a new trial. The final line of the displayFunc renders the new scene.

##### vi. Collect and save data

Researchers will likely want to specify which data to record and at what rates throughout a single trial of the experiment. In Ouvrai this is best done by defining two ancillary functions:

- **handleFrameData**: This function accumulates sampled data for each trial, where the individual samples are rendered “frames”. It is explicitly called within stateFunc, in FSM states during which data should be recorded (Supp. Fig. 2). The data typically include a timestamp for each frame, the current state of the FSM, and the current positions and rotations of scene objects of interest, particularly those that are controlled by input sources, such as the VR controllers (Supp. Fig. 1).
- **handleStateChange**: This function is provided to the FSM at initialization (see above) and called internally by the FSM to accumulate data related to state transitions. It minimally includes a timestamp and the new state, as shown in Supp. Fig. 1, but can also include other data. For example, in VR experiments, it can include the position and orientation of the camera object (i.e., the VR headset), which we might want to sample at a lower rate than the controllers.

On each trial, data is collected in a trial data object (Supp. Fig. 1). At the start of each trial, the trial data object is populated with parameter values that determine how the trial will proceed, such as target position and trial type. These values were initialized when the trial sequence was created, and at this point they are copied into the trial data object (Supp. Fig. 2, ‘SETUP’ state). The trial data object is also re-initialized with empty arrays for sampled data at the start of each trial. Sampled data includes values such as controller position and FSM state changes. While a trial is running, the two functions above are called within the experiment loop to append new samples to the arrays, including aligned arrays of timestamps. Note that frames and state changes are sampled at different rates, so each data type must have its own timestamp array (Supp. Fig. 1). Any other data generated during a trial that the researcher wishes to save, such as reaction times, movement durations, or other sampled data, can be manually added to the trial data object. This data can be added at any time between trial setup (in the ‘SETUP’ state) and the call at the end of the trial (in the ‘FINISH’ state) that saves the contents of the trial data object to Firebase. The FSM also includes a state that monitors whether the trial data was successfully saved to Firebase (i.e., the ‘ADVANCE’ state). If successful, the FSM will advance to the next trial. Following the last trial of the experiment, any new information that has been added to the experiment configuration object is saved to Firebase (note that changed values will not overwrite initial values that were saved after consent). Finally, Ouvrai makes a record in Firebase that links the participant’s crowdsourcing platform ID to their complete study data (Supp. Fig. 1).

#### C. Extensibility

Because it is based on JavaScript, Ouvrai also allows researchers to import classes and functions from packages that are third-party dependencies installed via Node Package Manager (npm). Researchers therefore have the option to incorporate a wide variety of new functionality into their studies, thanks to the large JavaScript developer community who make powerful packages freely available on npm. This means that Ouvrai studies are highly extensible, as the structure of the experiment loop and the design of the components place minimal constraints on what the researcher can do. Ouvrai comes with a core set of third-party dependencies, including Three.js for graphics and VR support, Tween.js for animations, and d3-array for array manipulations. Researchers can import classes and functions from these dependencies in their experiments without installing anything else, because Ouvrai itself is installed by default in all Ouvrai experiments. Other packages can be installed into individual experiments by navigating to the relevant experiment directory at the command line, and running npm’s *install* command.

## Experimental Results

To demonstrate the feasibility and data quality of remote VR studies, we used Ouvrai to create and run three studies based on standard motor learning experiments in which participants adapt to visuomotor rotations. The three tasks were chosen in part because they are known to produce distinctive, recognizable patterns of results in laboratory studies, described for each experiment below. As a result, it is best to evaluate our central claims about replicability and data quality simply by examining whether the data plotted in Figs. 2 to 5 (individual data and descriptive statistics for each group) conform to previous findings, rather than resorting to null-hypothesis significance tests. All experiments required participants to make reaching movements from a home position to one or more targets, while holding the right VR controller in their right hand. An example of a three-dimensional reach trajectory (out to a target and back to the home position) is shown in Fig. 2b. The unfiltered first-difference speed profile for this trajectory demonstrates the quality of the kinematic and timing data (Fig. 2c). The sample period was 8.41*±*0.84 ms (mean *±* SD), giving an average sample rate of 119 Hz

### Experiment 1: Spontaneous recovery

In Experiment 1, we establish that spontaneous recovery ^22, 23^ can be elicited and reliably measured in remote VR studies. Spontaneous recovery is a paradigm in which the motor memory of a previously learned perturbation (*P* ^+^; in this case a visuomotor rotation) is re-expressed after adaptation has been reduced to baseline by a brief intermediate exposure to the opposite perturbation (*P* ^+^). In Experiment 1, participants performed 120 reaching movements across four phases, using a pointer tool to hit a target ring that appeared straight in front of them (Fig. 2bd). Note this is considerably shorter than previous spontaneous recovery experiments, which lasted for 420–720 trials. However, this was found to be sufficient to elicit the recovery effect.

As shown in Fig. 3, individual reach angles (Fig. 3a) and group average reach angles (Fig. 3b) were consistently directed straight ahead during the baseline phase. Next, a clockwise visuomotor rotation of 15*^◦^* was introduced and participants adapted to this rotation for 60 trials (*P* ^+^ phase). In Fig. 3, the rotation magnitude is indicated with dashed lines showing the reach angle required to fully compensate for the rotation on each trial. Both the example individual (Fig. 3a) and the group average (Fig. 3b) exhibit a stereotypical learning curve for adaptation to an abruptly introduced visuomotor rotation, as reported in many laboratory studies. After the *P* ^+^ phase, the visuomotor rotation switched directions to 15*^◦^* counterclockwise for the next 10 trials (*P^−^* phase). By the end of the *P^−^* phase, adaptation to the *P* ^+^ appears to have been abolished; the average reach angle is approximately 5*^◦^* in the opposite direction of the initial adaptation (Fig. 3b).

**Fig 3.**
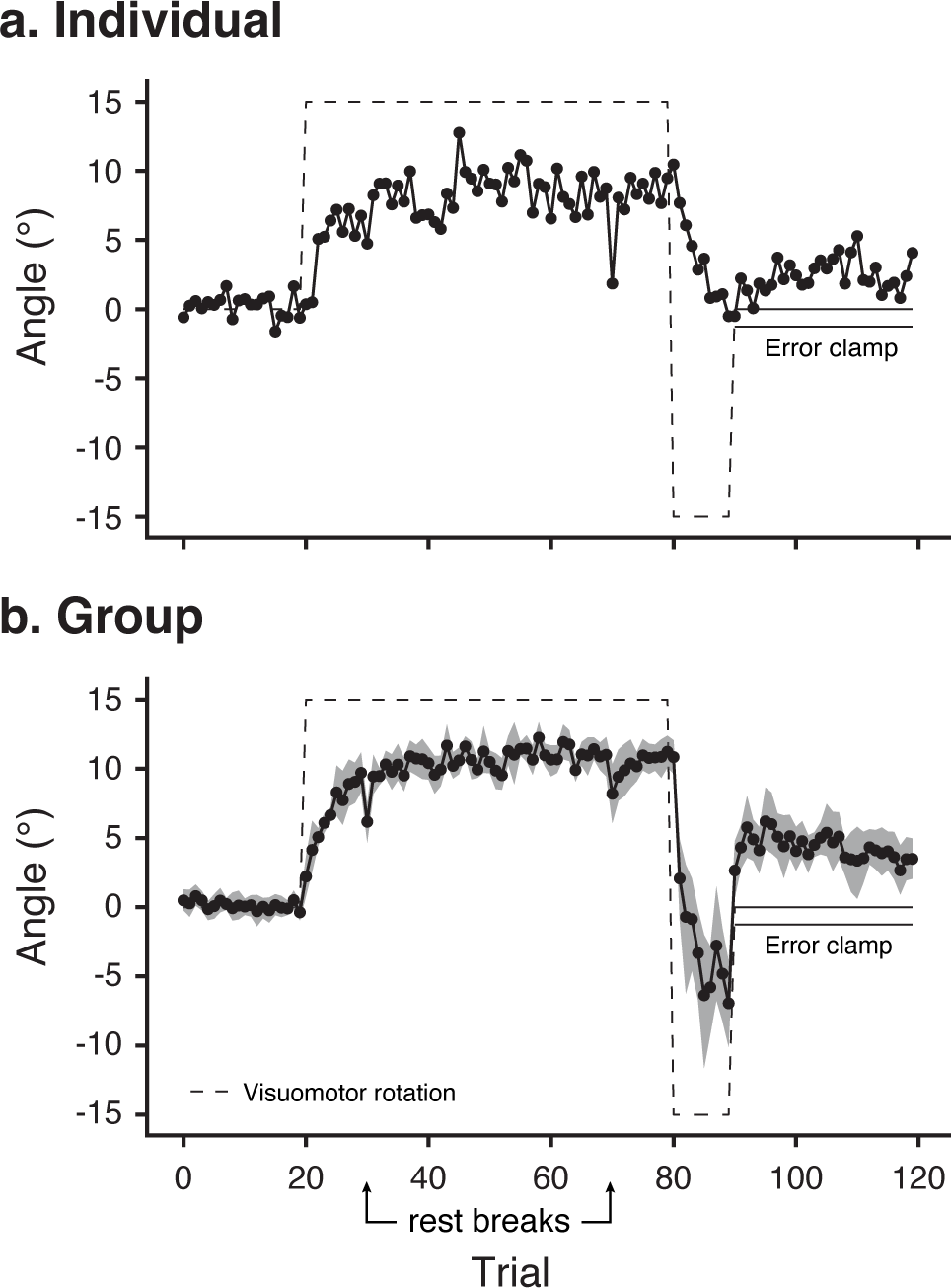
Experiment 1: Spontaneous recovery. The dashed line shows the reach angle that would be required to fully compensate for the visuomotor rotation. The final 30 trials are error-clamp trials (indicated by double lines). (a) Individual participant data. (b) Group average data. Shaded area shows bootstrapped 95% confidence intervals.

Crucially, however, in a final block of 30 trials, participants experienced no movement errors due to the introduction of an “error clamp” (see Methods). In the error-clamp phase, we found that a memory trace of the *P* ^+^ phase still remains, as evidenced by a progressively positive reach angle, which peaks in the third trial of this phase then slowly decays. This is the stereotypical profile of spontaneous recovery demonstrated previously in laboratory experiments, clearly visible in data from 10 participants performing an online unmoderated VR study. By replicating and precisely measuring the phenomenon, we demonstrate that Ouvrai could be used to efficiently examine variants of the basic spontaneous recovery paradigm in order to evaluate different theories of motor learning.

### Experiment 2: Dual adaptation with control points

In Experiment 2, we establish that online, unmoderated VR studies can be used to implement a second important feature of motor learning. This is the cue-dependent ability to simultaneously adapt to two opposing perturbations (for example, visuomotor rotations). In this experiment, each perturbation is associated with different control points on a hand-held tool ^24^. Participants made reaching movements to one of two targets with a T-shaped tool. The tool had red and blue control points on the left and right ends of the crossbar, respectively (see Fig. 2a). The targets were directly ahead of the associated control point and color coded to match it. During the baseline phase, no rotation was applied. Fig. 4 shows that reach angles for each control point tended to be indistinguishable during baseline phase, for both an individual participant (Fig. 4a) and in the group average (Fig. 4b).

**Fig 4.**
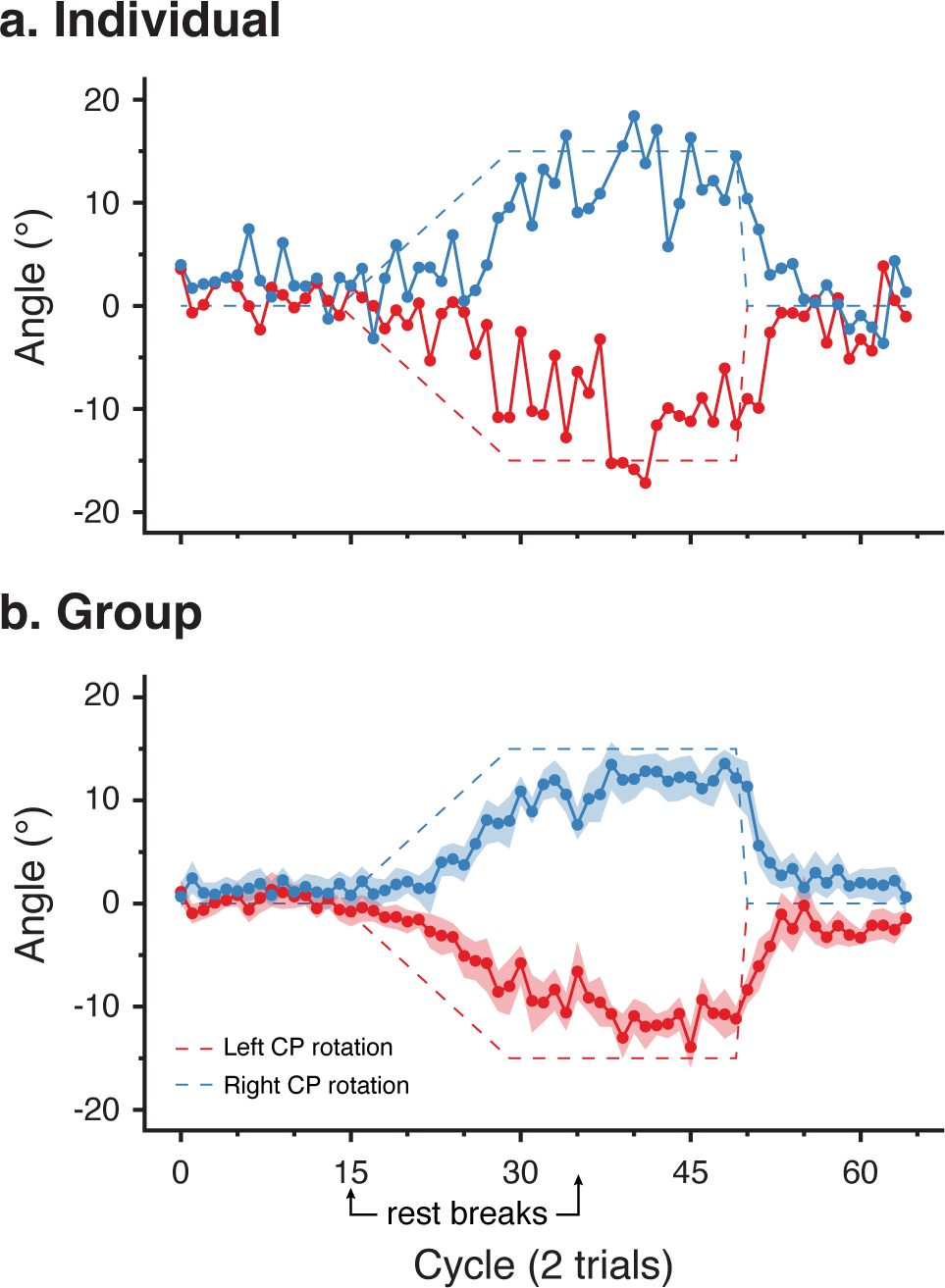
Experiment 2: Dual adaptation with control points. (plotted as in Figure 3). (a) Individual participant data. (b) Group average data. Shaded area shows bootstrapped 95% confidence intervals.

After 15 trial cycles (30 trials) of baseline reaching, the opposing visuomotor rotations associated with each control point were linearly ramped up. When reaching with the blue control point to the blue target, visual feedback of the tool was rotated clockwise. When reaching with the red control point, visual feedback was rotated counterclockwise. The rotations increased in magnitude by 1*^◦^* per trial cycle and plateaued at *±*15*^◦^* for 20 trial cycles. Fig. 4b shows that during the ramping phase, participants begin to aim on average to the left (positive reach angle) when using the blue control point and to the right (negative reach angle) when using the red control point. In the individual data, displayed in Fig. 4a, we observe that switching between the two perturbations caused some interference between the two motor memories, as evidenced by the increased variance in reach angle during the adaptation phase compared to the baseline phase. By the end of the plateau phase, all participants consistently reached leftward with the blue control point and rightward with the red control point.

Finally, in the de-adaptation phase with no rotations (as with the baseline phase), reaches were directed in different directions for each control point, separated by an angle that is greater than observed during baseline for at least the first 5 trial cycles. This is observed even for individual participants (e.g. Fig. 4a) and is evidence that the slow decay observed in the group average reflects gradual, implicit de-adaptation, and is not an artifact of averaging fast, explicit de-adaptation (i.e., suddenly reverting to the baseline aiming strategy) that occurs at different times across participants. These results demonstrate that a laboratory motor learning paradigm can be replicated using a remote VR study.

### Experiment 3: 3D generalization of visuomotor learning

In Experiment 3, we conducted a novel experiment that would take advantage of the 3D workspace available in VR. This is one of the qualitatively unique features of remote VR studies compared to studies that use a mouse, trackpad, or touchscreen. To this end, we adapted participants to a 16*^◦^*visuomotor rotation (in the horizontal plane, as in Experiments 1 and 2) experienced during reaches toward a single training target. Participants had to touch the training target with the tip of the pointer tool and press the trigger button to complete the reach. They received continuous visual feedback of the pointer during these reaches. However, in a series of no-feedback test trials before and after the adaptation phase, participants reached without visual feedback to 12 generalization targets distributed around the training target, *±*4 cm and *±*8 cm along each cardinal axis, as well as the training target. On these trials, since they received no visual feedback, participants pressed the trigger button when they believed that the tip of the pointer tool was inside the displayed target. The decision to press the button on these trials was therefore based on motor predictions and proprioceptive signals. These reaches allowed us to measure how the adaptation to the training target generalized over 3D space.

All participants showed consistent rightward biases in their reaching during the post-exposure generalization phase compared to the pre-exposure phase, showing broad generalization of adaptation over 3D space. This is shown for an example participant in Fig. 5a and in the group average in Fig. 5b. The pattern of biases in the group average data shows a rotational structure that qualitatively matches the imposed visuomotor rotation. This demonstrates that it is possible to successfully examine motor learning in three dimensions using an online unmoderated VR study.

**Fig 5.**
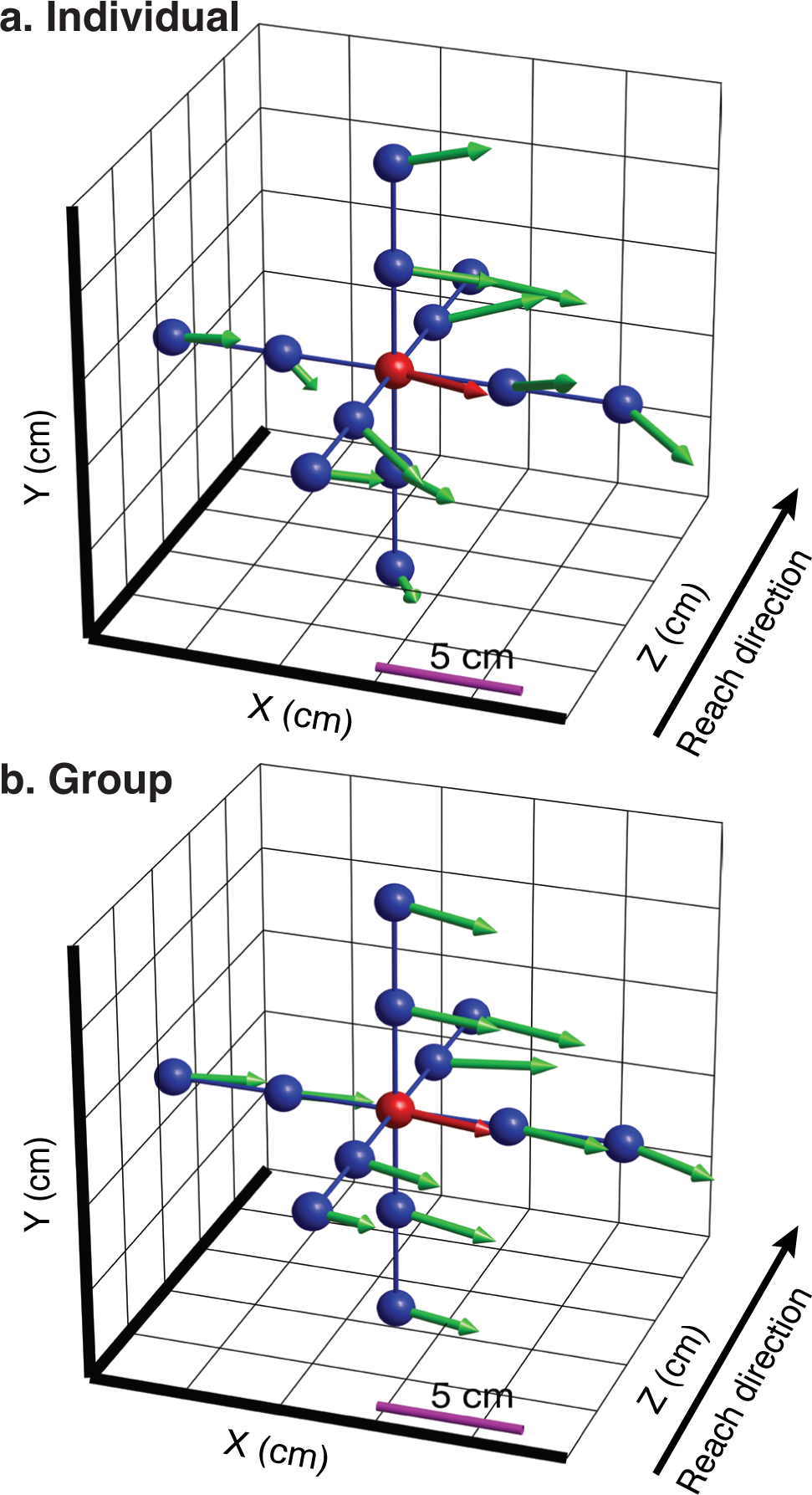
Experiment 3: 3D generalization of visuomotor learning. Participants adapted to a 16*^◦^* rotation at the central training target (red; the rotation leads to a 5.7-cm mismatch between visual and actual hand location at the target). They also performed no-feedback reaches to untrained generalization targets (blue) as well as the central target. Pre-exposure reach endpoints (not depicted) showed no systematic bias away from the displayed targets. The vectors show the shifted reach endpoints for each target in the post-exposure phase. (a) Individual participant data. (b) Group average data reflects extended spatial generalization of the learned rotation around the home position.

## Discussion

Virtual reality (VR) gives neuroscientists the ability to create immersive 3D environments and record the movements of human participants at high spatial and temporal resolution, enabling innovative studies of sensorimotor and cognitive functions ^25–27^. To date, nearly all VR studies have been moderated laboratory studies, which can be slow and costly. Crowdsourced research has emerged as a way to bypass the data collection bottleneck, but the possibility of conducting crowdsourced VR studies has not been realized due to the significant technical and software engineering challenges involved. We developed Ouvrai to address the need for open-source software that makes it easy for researchers to create, run, and analyze remote VR studies.

The major design goals for Ouvrai (described above) guided our choices about study infrastructure, the layout of the main experiment file, and the component library. We demonstrated that remote VR studies built with Ouvrai produce high-quality data, eliciting and reliably measuring effects that are important to current theoretical debates about the computational and neural underpinnings of motor learning. The results of our three validation studies (spontaneous recovery, dual adaptation, and 3D generalization) show that it is possible to parallelize data collection in experiments with immersion, interactivity, and data quality rivaling what is currently achievable in a physical laboratory, while substantially increasing data acquisition bandwidth and reducing costs.

Most importantly, the entire Ouvrai project is free and openly available for researchers to download and start using today. Interested researchers should consult the official documentation at https://ouvrai.com, which provides a practical walkthrough of how to create and run experiments with Ouvrai. In the remainder of the Discussion, we discuss the potential of Ouvrai in a broader research context, calling attention to features that we have glossed over or only briefly mentioned up to this point.

### Hand-tracking and eye/face-tracking

The latest generation of VR headsets (such as the widely owned Meta Quest 2 and newer models) provide hand tracking capabilities that use the externally-facing cameras on the headset to estimate the positions and orientations of 25 rigid bodies on each hand (corresponding to 19 joints). This allows researchers to perform remote VR studies even with populations of individuals that are unwilling or unable to hold VR controllers for the duration of the study, for instance younger children or individuals with diminished grip strength. Researchers can also collect data on individual finger movements, enabling studies on topics such as grip pre-shaping during grasping ^28, 29^, finger movement sequencing ^13, 30^, *de novo* learning of gesture-based controls ^31, 32^, and even body schema modification by virtual augmentation of the hand ^33^. Ouvrai comes with template code for a demo experiment that displays articulated 3D models of the two hands and shows researchers how to access and manipulate the hand-tracking data within an Ouvrai experiment. In this demo, the index fingers of the left and right hands are switched at the knuckle (as depicted in Fig. 2e), and the appearance of the hands can be changed from a skinned mesh to a set of 25 oriented cubes by crossing and uncrossing the wrists.

The latest generation of VR headsets include cameras inside the headset that can be used to recognize the wearer’s facial expressions, detect movements of the eyes, and estimate where the user is looking. Unfortunately, this data is not yet accessible in web VR applications because the group that creates the rules governing how VR support must be implemented in web browsers has not published specifications for eye-tracking or face-tracking APIs. This is due to concerns about the privacy ramifications of making such data available over the web (i.e., fingerprinting and profiling), as revealed by meeting notes ^34^ and GitHub discussions on this issue ^35^. Some progress has been made in that one member of the WebXR working group has created a draft of an expression-tracking API that relies on data from the inward-facing cameras, but this API explicitly avoids giving precise information on where the user is looking ^36^. As it is unclear what form the eye-tracking API(s) will eventually take (specifically, what level of abstraction will be imposed on the available data), we cannot predict whether high-precision studies of eye movements will be feasible. We expect further developments as more headsets supporting these features are sold, so interested researchers should follow this topic. Ouvrai will be updated to provide support and examples of how to use these features as soon as they become available.

### Can I run a non-VR study?

Ouvrai was designed specifically to enable remote VR studies, but the same workflow can be used to develop studies for computers and mobile devices (e.g., phones and tablets). Although these display devices have 2D screens, researchers have the option to present an orthogonal projection of a 2D workspace or a perspective projection of a 3D scene. Recent studies have shown that non-VR interfaces (i.e., 2D screens with mouse/trackpad and keyboard inputs) are sufficient to replicate laboratory results for some motor learning paradigms ^5, 6, 8, 9, 12, 13^. Additionally, although we recommend VR for greater immersion and improved data quality, there may be instances where a non-VR task is more appropriate for the aims of the study. For example, some recent online studies ^10, 11^ involved recruiting very large (N > 1000) and diverse samples, and it is unlikely that the sample sizes and diversity achieved in these studies could be matched with online VR studies.

To help researchers get started with non-VR studies, Ouvrai includes two non-VR template experiments (in addition to the four VR templates). These involve recording and responding to keyboard, trackpad, mouse, and/or touchscreen inputs. They provide all of the basic functionality that would be needed to implement a variety of motor tasks, such as the serial reaction time task where keyboard button presses are recorded to study motor sequencing ^13^, a visuomotor rotation paradigm where mouse/trackpad movements are recorded to study motor adaptation ^8–10^, or virtual physics tasks where the timing and/or location of clicks and button presses can be used to study learning of the mechanical properties of objects ^5–7^. Additionally, the touchscreen template experiment imports the *cannon-es* physics engine in order to simulate realistic 3D collisions, showcasing the extensibility of Ouvrai studies via JavaScript modules.

In VR studies, Ouvrai provides the XRInterface component to make it easy to present interactive UI elements in VR studies. Similarly there are components that implement helpful UI elements in non-VR studies, including the CSS2D, Fullscreen, Points, PointerLock, and Progressbar components. Unlike many existing crowdsourced research tools, Ouvrai does not focus on support for surveys. If you seek to run a simple survey, there are better tools than Ouvrai for this purpose. However, researchers who want to include a survey within an interactive Ouvrai study should note that the component library does include a customizable Survey component, typically to collect demographic information (e.g., handedness, device specifications, VR/gaming experience) before or after the primary task (see template experiments).

### Why not Unity?

When creating Ouvrai, we considered relying on the game development engine Unity, which is currently the most popular software for native (non-web) VR development. However, we found that Unity was not a viable solution for online studies. The documentation states plainly that “Unity does not support XR on WebGL” ^37^. There is a third-party package called WebXR Export that makes it possible to get a Unity VR app running on the web, but we found that it introduced several new problems. First, it was very complicated to correctly add JavaScript code to the resulting Unity app, making it hard to integrate with cloud services like Firebase, or to customize the appearance of the web page that displays the app. Second, it was not possible to test changes to the experiment code without compiling the whole project, which typically took several minutes, dramatically slowing iteration cycles during development. Lastly, the most attractive features of VR development with Unity, such as the built-in XR Management system and the Meta Quest SDKs that provide access to the latest headset features, were not available when using WebXR Export. Other drawbacks to using Unity include the fact that scripting must be done in C# and the tension between scripting and using the graphical editor. Furthermore, Unity cannot be used for free in an organization that has received revenue or funding in excess of $100K during the past 12 months. Overall, Unity does not easily integrate with web development tools and workflows. In contrast, 3D engines written specifically for the web platform, like Three.js, are focused on efficiently packaging and distributing code over the internet. Additionally, our decision to build Ouvrai around Three.js is consistent with the latest best-practices advice from Meta for developing web-based VR apps ^38^, so it is reasonable to expect support for this approach to continue into the future.

### Telerehabilitation and clinical research

One of the most promising potential avenues for Ouvrai to make an impact is in the realm of telerehabilitation and clinical research. A central aim of motor learning research in healthy human participants is to enable clinicians to design more effective rehabilitation protocols for patients with disorders of movement or coordination. Although we have focused on the utility of Ouvrai for expanding and accelerating research on healthy sensorimotor function, it is also an ideal platform for the development and deployment of telerehabilitation, the remote administration of rehabilitative treatment^39, 40^. VR-based telerehabilitation could provide personalized assessment and rehabilitation that optimizes motor learning and adapts to changes in individuals over time. To better understand the relative advantages and limitations of VR-based telerehabilitation, more data are needed, and clinician-scientists must build familiarity with conducting remote VR studies. By enabling researchers to quickly incorporate remote VR studies into their research programs, Ouvrai can help to overcome these key obstacles to progress in telerehabilitation. Patient-facing applications would particularly benefit from two core features: (1) immediate, secure transmission of movement data between client and database (i.e., patient and clinic), and (2) low-friction deployment of new VR experiences and updates to the patient. For clinical work, researchers should be sure to maintain HIPAA compliance (see https://cloud.google.com/security/compliance/hipaa for more details).

Likewise, many cognitive neuroscientists conduct clinical research on participants who have been diagnosed with a neurological disorder. Some of these disorders, like cerebellar degeneration, are quite rare and it can be expensive to bring patients into the laboratory. Ouvrai opens the door to an alternative approach in which a VR device can be provided to the patient for use in their own home. During an initial in-person visit (or, if necessary, a remotely conducted orientation), the researcher would demonstrate how the patient can use the headset to access the laboratory’s studies at their own convenience, and make sure they can do so safely. Thereafter, it becomes quite convenient to conduct a wide variety of studies, including longitudinal studies with higher session frequencies.

In summary, Ouvrai provides a new and alternative route to high-quality data that can complement laboratory studies. It can be used to pilot studies before they are performed in the laboratory or it can replace laboratory studies.

## Methods

All experiments required participants to make reaching movements from a home position to one or more targets, while holding the right VR controller in their right hand. The controller was visually rendered as a tool in VR. Before the experiment started, participants were instructed to sit up straight or stand, and to notice the tool in their right hand. They were instructed that they would use the tool to reach toward different targets. In order to calibrate the movement workspace to different participant heights, participants were required to hold the controller at chest height and press the trigger button.

Participants were instructed about the number of trials they would perform and the number of enforced rest breaks. They were also given the option to rest at any time by not returning to the home position after a successful reach. During enforced rest breaks (which were a minimum of 15 seconds before participants could resume the experiment), participants were instructed to take a short break, stretch and relax their arm, and to not exit or remove their headset.

When conducting moderated experiments in the lab, the researcher has the opportunity to explain and demonstrate the task to the participant, and then to provide verbal feedback while the participant performs some number of practice trials. Since Ouvrai studies are online and unmoderated, this process must be automated and included as part of the early stages of the experiment. In our case, before starting the experiment, we showed participants a “demo avatar” that was a darker version of the same tool held by the participant. We animated the position of the demo avatar so that it made repeated out-and-back reaching movements (with some randomness in the endpoint of each reach), similar in direction and magnitude to the movements the participant would be required to make. Instructions were displayed which asked participants whether they could comfortably perform the movements, along with suggestions on how to obtain a more comfortable pose with respect to the targets: physically move their body, recenter their view using a button on the VR headset, or reset the chest height calibration.

Finally, the first trial of all experiments was always a “practice trial” in which participants were provided with step-by-step instruction text guiding them through a single trial. For example, at the start of a practice trial, participants were instructed to place the end of the tool in the home position cube, noting that the cube would turn black when they were correctly positioned. When the target appeared, participants were instructed to reach towards it, and after hitting the target, they were instructed to return to the home position. When they completed the practice trial, they were asked whether they were ready to start, or if they would like to repeat the trial and review the step-by-step instructions. Participants could complete up to three practice trials.

### Participants

Using the Prolific crowdsourcing platform, we recruited 30 participants (23 males, 7 females) between 18 and 57 years old (median 59) and residing in 14 different countries. Participants were of four ethnicities (24 White, 4 Mixed, 1 Asian, 1 Other) and their prior experience with Prolific varied widely, ranging from 6 to 3627 approved studies (median 275). Participants in Experiments 1 and 2 were paid $5, and participants in Experiment 3 were paid $6. We used the following Prolific screening criteria: approval rate of at least 95%, age 18–65, fluent in English, normal or corrected-to-normal vision, VR headset ownership, and no diagnosis of multiple sclerosis, mild cognitive impairment, dementia, or of any other mental illness that is uncontrolled and has a significant impact on daily life. All experiments were conducted in accordance with the 1964 Declaration of Helsinki, following protocols approved by the Columbia University Institutional Review Board. Informed consent was obtained from all participants prior to their participation.

### Experiment 1: Spontaneous recovery

Spontaneous recovery refers to a motor learning paradigm in which participants’ movements are initially adapted to a single perturbation (*P* ^+^), typically a curl field or visuomotor rotation, and then de-adapted by brief exposure to an opposing perturbation (*P^−^*), a curl field or visuomotor rotation with the same magnitude but opposite direction as *P* ^+^. Participants then perform a series of “error-clamp” trials in which error feedback is suppressed. During these trials, participants quickly revert to making movements that would partially compensate for the initial perturbation (*P* ^+^). This re-expression of the earlier motor memory, which had been behaviorally extinguished by the conflicting perturbation (*P^−^*), is known as spontaneous recovery ^22, 41^.

The phenomenon of spontaneous recovery was originally explained by a model of motor learning consisting of two motor memories, which adapt and decay at different rates, summing their outputs to give the net adaptive state (the dual-rate model ^22^). Subsequently, spontaneous recovery provided evidence for a model based on contextual inference, which allows for the creation of new motor memories and the recall of existing memories (the COIN model ^41^).

In Experiment 1, we implemented a spontaneous-recovery paradigm using opposing visuomotor rotations in a remote VR study. Participants made reaching movements from a home position to a target. On each trial, once participants were at the home position, the target appeared after a 200-ms delay. When they hit the target, participants were given visual feedback (a 220-ms animation of the target expanding then shrinking until it disappeared), audio feedback (a “pop” sound) and haptic feedback (a 40-ms controller vibration at 60% intensity). Neither reaction times nor movement times were constrained.

The home position was a 1.5-cm wire cube and the target was a torus (major radius 2 cm in the vertical plane, minor radius 0.2 cm). The controller location was displayed as a cylindrical metallic gray rod (9 cm long, radius 1 cm) with a white spherical tip, resembling a small pointer or wand (Fig. 2d). Participants reached with the pointer tool to a target ring located 20 cm straight ahead.

Participants performed 120 reaches with 20 baseline trials, 60 *P* ^+^ adaptation trials, 10 *P^−^* de-adaptation trials, and 30 error-clamp trials. *P* ^+^ and *P^−^* were visuomotor rotations of +15*^◦^*and -15*^◦^*, respectively. Terminal-feedback trials began after trial 12 for the remainder of the experiment. In these trials, visual feedback of the tool was withheld while it was 3 to 17 cm from the home position. This no-feedback region was shown as a transparent gray area, in which an expanding black ring showed their distance towards the target but not the angle. The tool reappeared at the end of the reach (see Fig. 2a for an example of a terminal-feedback trial, in this case for Experiment 2). Participants were shown the following instructions before the first terminal-feedback trial: “Try to hit the target without visual feedback! In the gray area, the tool will disappear. A dark ring shows your distance.” On error-clamp trials, the terminal feedback was always shown in the correct direction to reach the target, independent of where the participant reached. They were explicitly notified of this error clamp and instructed to reach straight towards the target, that is, to stop intentionally reaching to either side of the target if they had adopted such a strategy. Enforced rest breaks were given after trials 30 and 70.

### Experiment 2: Dual adaptation with control points

Although participants easily adapt to single dynamic or visuomotor perturbations, when two opposing perturbations are presented sequentially, interference is observed and participants fail to adapt ^42–46^. However, when each perturbation is paired with a different sensorimotor cue, participants can adapt to opposing perturbations ^47–50^. In this case, a motor memory for each perturbation becomes associated with its paired cue. A range of different cues for adaptation to opposing perturbations have been explored. Moreover, interference paradigms have been used to examine a number of questions in motor control, including the representation of different movement types ^47, 49^, the representation of different perturbations ^46^, the consolidation of motor memories ^43^, the time-dependent encoding of motor memories ^50^, motor planning ^51, 52^, and motor imagery ^53^.

Recently, an interference paradigm showed that opposing dynamic perturbations can be learned when each is associated with the control of a different point on a hand-held tool ^24^. In Experiment 2, we implemented a control-point paradigm using opposing visuomotor rotations in a remote VR study.

This experiment was similar to Experiment 1 so we only describe the differences. The tool was T-shaped, consisting of a cylindrical handle (9 cm long, radius 1 cm) and a cylindrical crossbar (16 cm long, radius 1 cm), with red and blue spherical tips at each end of the crossbar (Fig. 2a). These colored tips were the left and right control points of the tool. The handle of the T-shaped tool tracked the position and orientation of the handheld controller, whereas the crossbar had a fixed orientation (i.e., it would only translate, even as the handle it was attached to rotated). To begin a trial, participants had to position the tool so that the left and right control points were inside corresponding left and right home positions. Participants used the tool to make reaching movements to target rings located 20 cm straight ahead of the corresponding home positions. The targets were color coded to match the control point, with the order of the targets (left/red and right/blue) randomized within each pair of reaches. To complete a trial, participants were required to move the appropriate control point so that it passed through the target.

Participants performed 120 trials with 30 baseline trials, 60 adaptation trials (visuomotor rotations of -15*^◦^* for the left control point and +15*^◦^* for the right control point, with the rotation linearly ramped up over the first 30 trials), and 30 de-adaptation trials with no rotations. Terminal-feedback trials began after trial 14 for the remainder of the experiment. Enforced rest breaks were given after trials 30 and 70.

### Experiment 3: 3D generalization of visuomotor learning

A number of studies across different behavioral paradigms have shown that motor learning shows graded generalization ^44, 54–57^. For example, if participants are adapted to a visuomotor rotation at a single target, expression of the resulting adaptation at novel targets decreases as their distance from the trained target increases ^58^. Generalization is an important feature of motor learning which has revealed details of the computational and representational mechanisms in the motor system ^54, 56, 59, 60^. In Experiment 3, we implemented a generalization paradigm in 3D using a visuomotor rotation in a remote VR study that was similar to Experiment 1.

Participants used the tool as in Experiment 1 to make reaching movements from a home position to targets (textured spheres of radius 1.4 cm) which included a training target 20 cm directly ahead, and 12 generalization targets, spaced *±*4 cm and *±*8 cm on each of the 3 axes, relative to the training target (4 targets per axis to give the 12 generalization targets).

In total 180 trials were performed. Trials consisted of 26 full-feedback baseline trials to the exposure target (2 trials per target, no rotation), 26 pre-exposure no-feedback trials to all targets (2 trials per target, no rotation) interleaved with 26 full-feedback reaches to the exposure target, 50 full-feedback adaptation trials to the exposure target (a visuomotor rotation of 16*^◦^* was linearly ramped up over the first 20 trials), and 26 post-exposure no-feedback trials interleaved with 26 full-feedback trials, as in the pre-exposure phase.

On all trials, participants were required to press the trigger button on the controller when they had reached each target. On no-feedback trials, feedback of the tool was withheld when the tool left the home position. Participants were shown the following instructions before the first no-feedback trial: “Try to touch the targets without visual feedback! Sometimes the tool will disappear as you reach forward. Press the trigger when you think you are touching the target.” Enforced rest breaks were given after trials 20, 60, 100 and 140.

### Analysis

For each participant in the spontaneous-recovery and dual-adaptation experiments, we calculated the reach angle at a distance of 17 cm from the start on each trial (just before the tool reappeared in the no-feedback trials). One trial from one participant was excluded because a valid reach angle could not be extracted. We then computed the group average along with bootstrapped 95% confidence intervals (1000 samples).

In the generalization experiment, the tool position was recorded in the pre-exposure (baseline) and postexposure no-feedback trials when the participant pressed the trigger button (when they believed they were touching the target). During each phase, for each participant, we calculated the average position across the trials for each target. We calculated the group average using these participant averages.

---

**Supp. Fig. 1].**
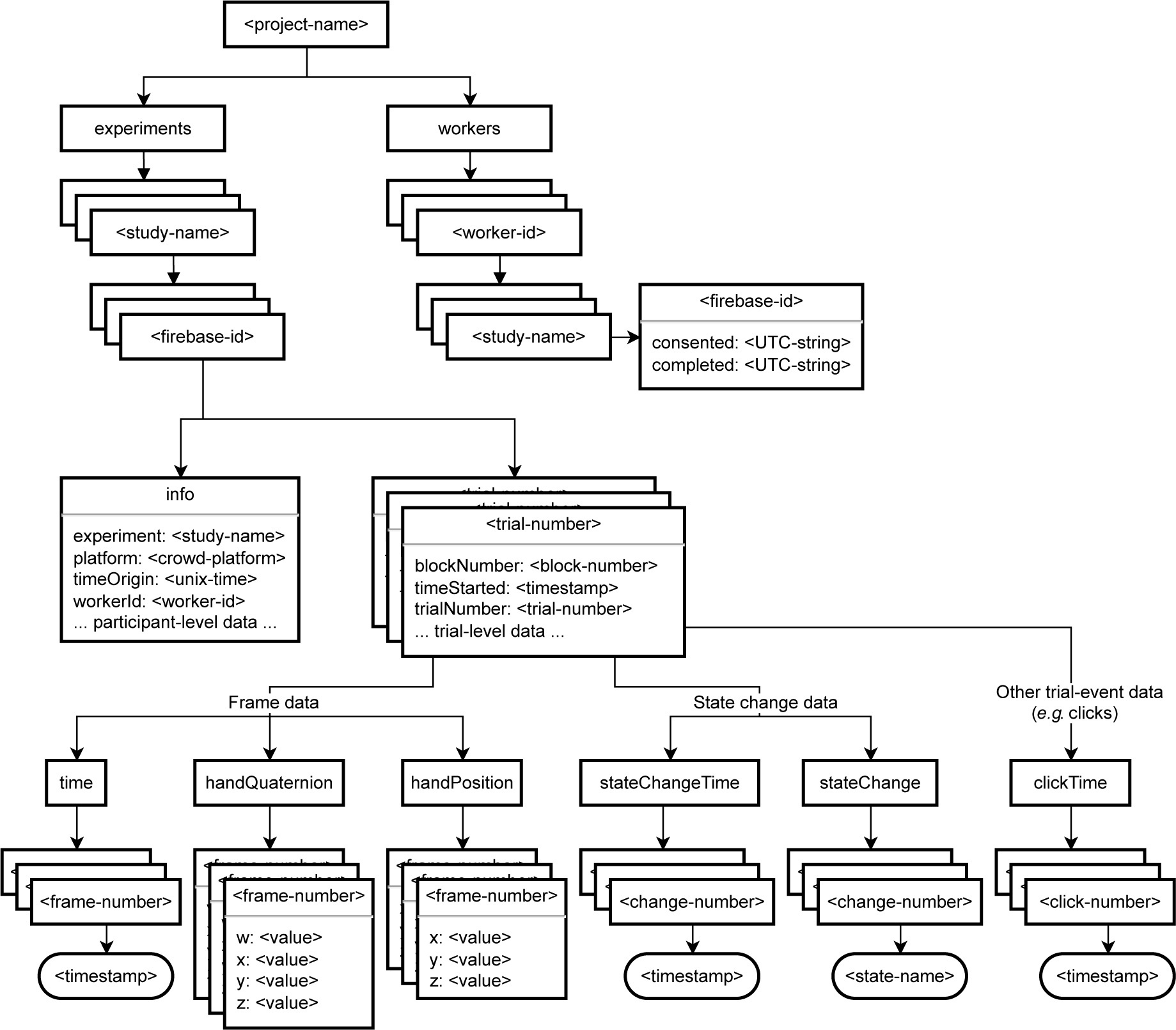
Layout of the Firebase Realtime Database. Data is structured hierarchically as a JavaScript object. An object contains one or more scalar-valued “fields” (typically strings, but sometimes integers), where each field points to a “value” which can be either a scalar value or a nested JavaScript object. The “root” of the database has a single field called *<project-name>* (angle brackets indicate variables) which is the name of the researcher’s Firebase project. The value of the *<project-name>* field is a nested object with two fields: “experiments” and “workers”. The value of the “workers” field is a nested object where the fields are the worker IDs (i.e., participant identifiers assigned by crowdsourcing platforms) of all participants who have ever participated in any of this researcher’s Ouvrai studies. The value of each *<worker-id>* field is a nested object where the fields are the names of all studies performed by this participant. The value of each *<study-name>* field is a nested object where the field(s) is the Firebase ID string that was assigned to this participant (discussed below). Note that multiple Firebase ID fields will appear here if the same crowdsourced participant completes the consent form of an Ouvrai study more than once. Finally, the value of the *<firebase-id>* field(s) is a nested object with two fields, “consented” and “completed”. These are “leaf” fields in the database (i.e., they have scalar values) containing timestamps indicating when the participant gave informed consent and when they completed the study (these timestamps are strings in Coordinated Universal Time). The primary purpose of the “workers” branch described above is to allow a researcher (or an Ouvrai utility) to find the data provided by a particular participant for a particular study, given only the worker ID assigned by the crowdsourcing platform. Experimental data is stored in the “experiments” branch of the database, under a particular Firebase ID in order to ensure database security. The value of the “experiments” field is a nested object with multiple fields that are the names of all studies for which this researcher has saved any data. The value of each *<study-name>* field is a nested object with multiple fields that are the Firebase IDs assigned for this study (i.e., anonymous IDs assigned by Firebase to individually identify visitors to the publicly hosted Ouvrai study page). Ouvrai uses the Anonymous Authentication feature of Firebase along with a custom set of database security rules to keep track of individual participants and maintain database security while preserving anonymity. Using this feature, each visitor to an Ouvrai study website who completes the consent form is assigned a temporary but unique ID (alphanumeric string, currently 28 characters), which cannot be “spoofed” by other users. After being assigned an ID, the participant (or rather, the Ouvrai study code that is running in their browser as they perform the study) can write data to the Firebase database in the “experiments” branch, but only at locations that respect the following path structure: /*<project-name>*/experiments/*<study-name>*/*<firebase-id>*/*<data-name>*, where *<data-name>* can be either “info” or an integer *<trial-number>*. Thus, the value of each *<firebase-id>* field is a nested object with multiple fields, one of which is “info” and the remaining are integer trial numbers, one for each trial in the experiment starting from zero. The value of the “info” field is a nested object with many different field-value pairs containing global experiment parameters. For instance, the field “experiment” has a string value that is the name of the Ouvrai study, and so on for “platform” (a string value that is the name of the crowdsourcing platform), “timeOrigin” (the UTC time in milliseconds at which the page was loaded), “workerId” (the crowdsourcing platform ID described above), and any other field-value pairs specified by the researcher in the experiment configuration object in the Ouvrai experiment code. The “info” object is saved at the time of consent and updated at the time of study completion. Experimental data is stored in the “experiments” branch of the database, under a particul Trial data is saved after each trial, producing consecutive, integer-valued *<trial-number>* fields. The value of each *<trial-number>* field is a nested object typically containing fields with scalar values describing properties of that trial (“blockNumber”, “trialNumber”, “timeStarted”) and typically others that are specific to the individual trials of a particular experiment (the location of a target object, the magnitude of a visuomotor perturbation, or the type of visual feedback provided). The trial data object also contains fields referring to data that was sampled at different rates during the trial. The values of these fields are nested objects with consecutive integer-valued fields indexing the individual samples (note this sampled data is actually stored in an array when it is saved in the Ouvrai experiment code). For example, in our VR studies, the controller data were sampled on each render frame to track reaching movements. This type of data is indicated by the descending arrows labeled “frame data”. One field related to frame data is called “time”, and its value is a nested object with multiple fields that are (as mentioned above) consecutive integer-valued frame numbers. The value of each *<frame-number>* field is a millisecond timestamp for the render frame, indicating when the sampled data (described next) was recorded. Two types of frame data that we typically collect during VR studies are denoted by the fields “handQuaternion” and “handPosition”. These contain the same set of *<frame-number>* fields as the “time” field, but the value of each *<frame-number>* field contains the relevant sampled data, i.e., a nested object with field-value pairs that represent the quaternion or position of the VR controller. Researchers may want to sample data at lower rates, for example only when the state of the finite state machine changes (see Supp. Fig. 2), or when the mouse is clicked. The remaining boxes in the lower-right illustrate how these trial events are sampled and incorporated into the database structure in a way similar to the frame data. Each type of event (i.e., each sampling rate) will typically include an array of timestamp data in addition to any other aligned arrays of recorded data.

**Supp. Fig. 2].**
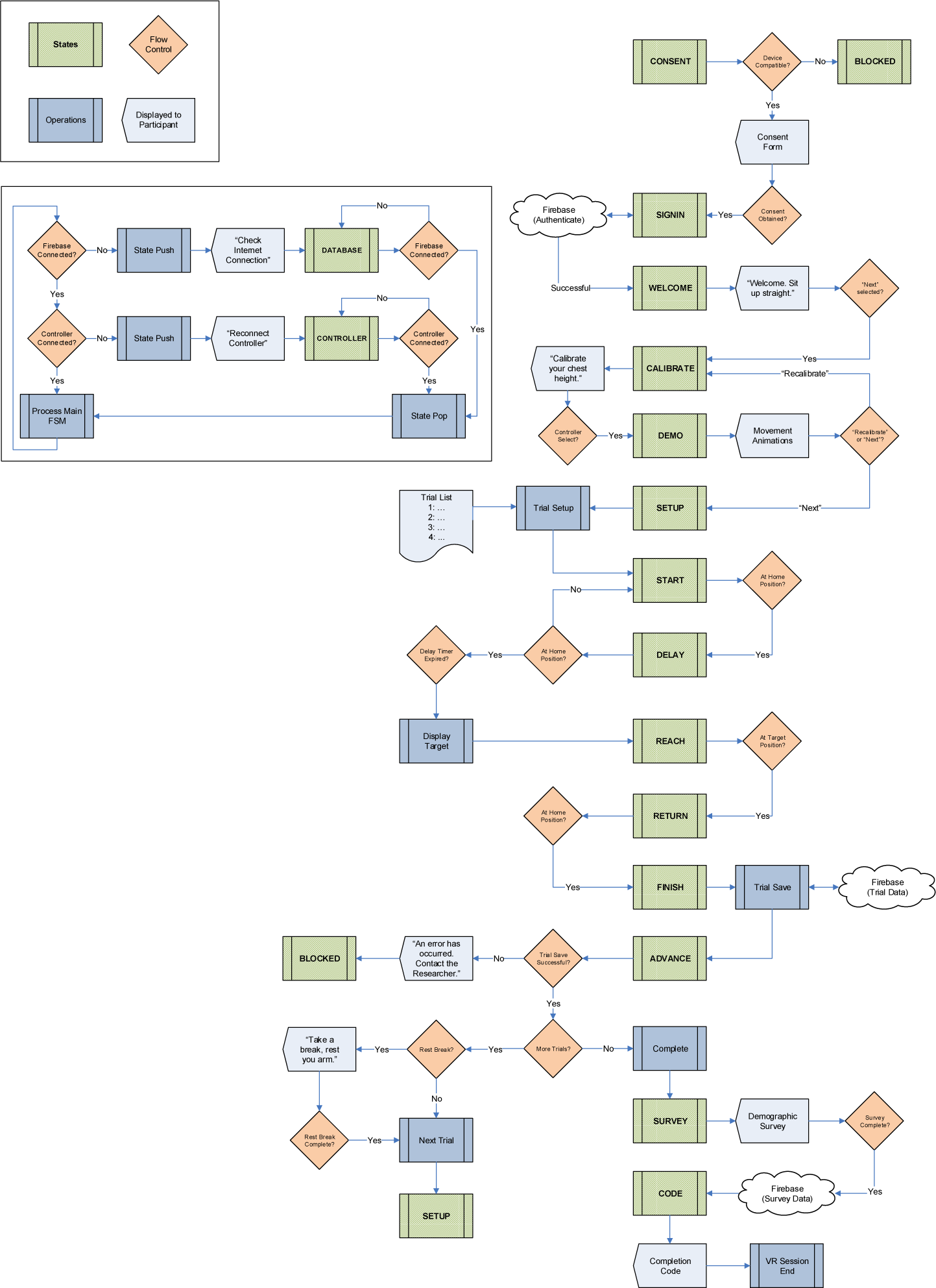
The finite state machine (FSM) for Experiment 1, represented as a flow chart. The key to symbols are shown in the top left inset. Green boxes represent states. Blue boxes represent operations or functions. Orange diamonds represent conditional flow control. Pale blue rectangular polygons represent text or other objects displayed to the participant (text can include input fields, such as surveys). Each state is associated with a specific action or phase of the experiment, where operations can be performed, flow-control conditions can be tested, and text or other objects can be displayed to participants. Flow control branches can test conditions of the run-time environment (the CONSENT state tests that the experiment is running on a compatible device), user input (the DEMO state waits for the user to select “Next” or “Recalibrate”), timers (the DELAY state waits for 200 ms), and experimental variables (the START state waits for the user to position the tool at the home position). The experiment proper, where participants are performing trials, is represented by states SETUP to ADVANCE. When states appear more than once (for example, the SETUP state), this is to avoid convoluted transition arrows. The larger inset (below the symbols key) represents interrupt states. These monitor critical conditions that must be resolved before the experiment can resume. For example, if the connection to Firebase is lost, the current state is saved (pushed), users are appropriately notified, and the FSM remains in the DATABASE state until the connection is re-established. Upon exiting an interrupt state, the saved state is restored (popped), and the experiment resumes. The inset also shows the iterative nature of the FSM, with the main branch represented as an operation (Process Main FSM). Note that some details have been omitted for brevity.

## Footnotes

i The constraint of running in the browser has shaped the kinds of studies that are run remotely, which in turn has influenced what scientists believe is achievable with an online study. The crowdsourced research landscape is dominated by surveys and (in a distant second place) simple behavioral tasks that display text or images and record mouse clicks or keyboard button-presses. For the most part, features like 3D graphics, rich interactivity, first-person navigation, and unconstrained limb movements are not considered viable in browser-based tasks.

ii The MTurk fee increases to 40% if a researcher posts a study configured to recruit more than 9 participants. However, by default Ouvrai avoids this fee by posting multiple copies of the same study, each configured to recruit 9 or fewer participants. Normally, this would allow the same participant to complete and submit the study multiple times (in theory, as many times as copies of the study were posted). However, with Ouvrai, each time a prospective participant views the study, the MTurk study page runs a routine that checks for the participant’s MTurk ID in the database to determine if they have previously submitted the study and, if so, notifies the participant that they cannot complete the study multiple times and hides the study link and submit button.

## Notes

https://ouvrai.com

https://github.com/EvanCesanek/Ouvrai

### Competing Interest Statement

The authors have declared no competing interest.

### Summary of Updates

Minor changes to formatting and references.

